# Turning the tables: Loss of adaptive immunity reverses sex differences in tuberculosis

**DOI:** 10.1101/2024.09.24.614709

**Authors:** David Hertz, Lars Eggers, Linda von Borstel, Torsten Goldmann, Hannelore Lotter, Bianca E. Schneider

**Affiliations:** Host determinants in lung infections, Priority Research Area Infections, Research Center Borstel, Leibniz Lung Center, Borstel, Germany; Core Facility Histology, Research Center Borstel, Leibniz Lung Center, Borstel, Germany; Airway Research Center North (ARCN), Member of the German Center for Lung Research (DZL), Großhansdorf, Germany; Department of Molecular Parasitology and Immunology, Bernhard Nocht Institute for Tropical Medicine, 20359 Hamburg, Germany

## Abstract

Sex-based differences in innate immunity may play a crucial role in susceptibility to and progression of tuberculosis (TB), a disease that disproportionately affects men. This study aimed to examine whether early host-pathogen interactions contribute to the heightened vulnerability of males to *Mycobacterium tuberculosis* (*Mtb*) infection. Using recombination activating gene knockout (RAG KO) mice, which lack adaptive immunity, we were able to isolate and analyze innate immune responses to *Mtb* without the influence of T and B cells. Surprisingly, and in stark contrast to wild-type mice that reflect the male bias as observed in humans, female RAG KO mice were more susceptible to *Mtb* than their male counterparts. Increased lung CFU in females was accompanied by a significant rise in inflammation, indicated by elevated levels of inflammatory cytokines and chemokines, as well as a massive influx of neutrophils into the lungs. In contrast, male mice exhibited higher levels of IFN-γ and CCL5, along with a greater presence of NK cells in their lungs, suggesting that, in the absence of adaptive immunity, males benefit from a more robust NK cell response, potentially offering greater protection by better controlling inflammation and slowing disease progression.

## Introduction

Tuberculosis (TB) is the most prevalent bacterial infectious disease in humans. The causative agent, *Mycobacterium tuberculosis* (*Mtb*), is carried by an estimated 2-3 billion people globally and claims approximately 1.5 million lives each year ^1^. Like many other infectious diseases, TB exhibits a strong male preponderance in disease development, with males being nearly twice as likely to have active TB compared to females. Historically, this difference has been attributed to behavioral factors such as higher rates of smoking and alcohol consumption among males ^2^. However, recent research, including our own, has begun to shed light on biological reasons behind the differing risks of TB between males and females ^3^. Particularly, studies using mouse models that isolate the impact of biological sex on TB susceptibility emphasize its importance, demonstrating that males experience a more rapid progression of TB and earlier death ^4-6^.

Increased male susceptibility to *Mtb* manifests in the chronic phase of the infection and is accompanied by marked differences in the organization of lesions in the lung ^5,6^. While the underlying reasons for these differences in lesion formation remain to be elucidated, it has become clear that the early stages of infection are crucial in determining its final outcome ^7^. Consequently, inherent functional differences among innate immune cells involved in early host-pathogen interaction could profoundly influence the overall outcome of the infection. Therefore, our aim was to investigate potential sex differences during early *Mtb* infection while excluding the influence of adaptive immunity. To do so, we studied recombination activating gene knockout (RAG KO) mice, which lack mature B and T lymphocytes ^8^. Unexpectedly, while males are typically more susceptible to *Mtb* in immunocompetent mice, RAG KO mice revealed the opposite pattern. Female RAG KO mice infected with *Mtb* HN878 exhibited higher bacterial loads, more severe weight loss, and earlier mortality compared to their male counterparts. The inflammatory response in females was more pronounced, marked by neutrophil-dominated lung pathology. In contrast, the increased presence of NK cells and significantly higher IFN-γ levels in the lungs of males suggest that they benefit from a more robust NK cell response, which may play a role in regulating inflammation and slowing disease progression. These findings highlight significant sex-specific differences in the innate immune response to *Mtb* infection, with females, unexpectedly, showing increased susceptibility to severe infection and inflammation when adaptive immunity is absent.

## Material and Methods

### Mice

C57BL/6j mice were bred under specific-pathogen-free (SPF) conditions at the Research Center Borstel. RAG KO mice were bred under SPF conditions at the Bernhard Nocht Institute for Tropical Medicine, Hamburg, Germany, and transferred to the experimental animal facility at the Research Center Borstel at least 5 weeks before the experiments for acclimatization. For infection experiments, male and female mice aged between 12-16 weeks were maintained under barrier conditions in the BSL 3 facility at the Research Center Borstel in individually ventilated cages. Animal experiments were in accordance with the German Animal Protection Law and approved by the Ethics Committee for Animal Experiments of the Ministry of Agriculture, Rural Areas, European Affairs and Consumer Protection of the State of Schleswig-Holstein.

### Mtb infection and determination of bacterial load

*Mtb* HN878 was grown in Middlebrook 7H9 broth (BD Biosciences, Franklin Lakes, USA) supplemented with 10% v/v OADC (Oleic acid, Albumin, Dextrose, Catalase) enrichment medium (BD Biosciences, Franklin Lakes, USA), 0,2% v/v Glycerol and 0,05% v/v Tween 80 to logarithmic growth phase (OD_600_ 0.2 - 0.4) and aliquots were frozen at -80°C. Viable cell number in thawed aliquots were determined by plating serial dilutions onto Middlebrook 7H11 agar plates supplemented with 10% v/v OADC and 0,5% v/v Glycerol followed by incubation at 37°C for 3-4 weeks. For infection of experimental animals, *Mtb* stocks were diluted in sterile distilled water at a concentration providing an uptake of approximately 100-200 viable bacilli per lung. Infection was performed via the respiratory route by using an aerosol chamber (Glas-Col, Terre-Haute, IN, USA) as described previously ^9^. The inoculum size was quantified 24 h after infection in the lungs and bacterial loads in lung, mediastinal lymph nodes, spleen and liver were evaluated at different time points after aerosol infection by determining colony forming units (CFU). Organs were removed ssaseptically, weighed, and homogenized in 0.05% v/v Tween 20 in PBS containing a proteinase inhibitor cocktail (Roche, Basel, Switzerland) prepared according to the manufacturer’s instructions for subsequent quantification of cytokines (see below). Tenfold serial dilutions of organ homogenates in sterile water/1% v/v Tween 80/1% w/v albumin were plated onto Middlebrook 7H11 agar plates supplemented with 10% v/v OADC and incubated at 37°C for 3–4 weeks.

### Clinical Score

Clinical score was used to indicate severity of disease progression. Animals were scored in terms of general behavior, activity, feeding habits, and weight gain or loss. Each of the criteria is assigned score points from 1 to 5 with 1 being the best and 5 the worst. The mean of the score points represents the overall score for an animal. Animals with severe symptoms (reaching a clinical score of ≥3.5) were euthanized to avoid unnecessary suffering, and the time point was recorded as the end point of survival for that individual mouse.

### Multiplex cytokine assay

The concentrations of various cytokines and chemokines in lung homogenates were determined by LEGENDplex™ (Mouse Inflammation panel and Mouse Proinflammatory Chemokine Panel, BioLegend) according to the manufacturer’s protocol.

### Histology

Superior lung lobes from infected mice were fixed with 4% w/v paraformaldehyde for 24 h, embedded in paraffin, and sectioned (4 μm). Tissue sections were stained with hematoxylin and eosin (HE) to assess overall granulomatous inflammation. Macrophages were detected by a polyclonal rabbit anti CD68 (abcam), neutrophils by monoclonal rat anti neutrophils (clone 7/4, Cedarlane Laboratories) and NK cells by a rabbit recombinant monoclonal anti NKR-P1C (EPR22990-31, abcam), respectively, followed by secondary antibody (biotinylated goat anti-rabbit or biotinylated goat anti-rat; Dianova), amplification (avidin-HRP) and color reaction (DAB solution; Vectastain). Slides were imaged with PhenoImager™ HT instrument (Akoya Biosciences). Affected lung area was quantified in relation to the whole lung area and the number of the different cell subsets in relation to all cells using the software QuPath (freeware).

### Statistical Analysis

All data were analyzed using GraphPad Prism 10 (GraphPad Software, La Jolla, USA). Statistical tests are indicated in the individual figure legends. Correlation was determined by calculating Pearson’s coefficient using a two-tailed analysis. A correlation was taken into account as of r≥ 0.60 (defined as strong correlation). Values of *p≤0.05, **p≤0.01, ***p≤0.001 and ****p≤0.0001 were considered significant.

## Results

### Female RAG KO mice are more susceptible to Mtb HN878 compared to males

To test the ability of male and female mice to control HN878 infection in the absence of adaptive immunity, we infected RAG KO mice and monitored disease progression and survival according to a clinical score, as described in the Materials and Methods. Moreover, we determined the bacterial load in the lungs, mediastinal lymph nodes (LN), spleen, and liver after 13 days, 21 days, at a flex time point, and in moribund animals that had reached the clinical endpoint according to our clinical scoring (Fig. 1A). The flex time points were chosen as follows: as soon as a mouse reached a score of 3, a randomly selected mouse of the opposite sex, which had been assigned before the experiment, was also taken, regardless of its score. This allowed us to determine the CFU in relation to the disease stage and progression.

**Figure 1.**
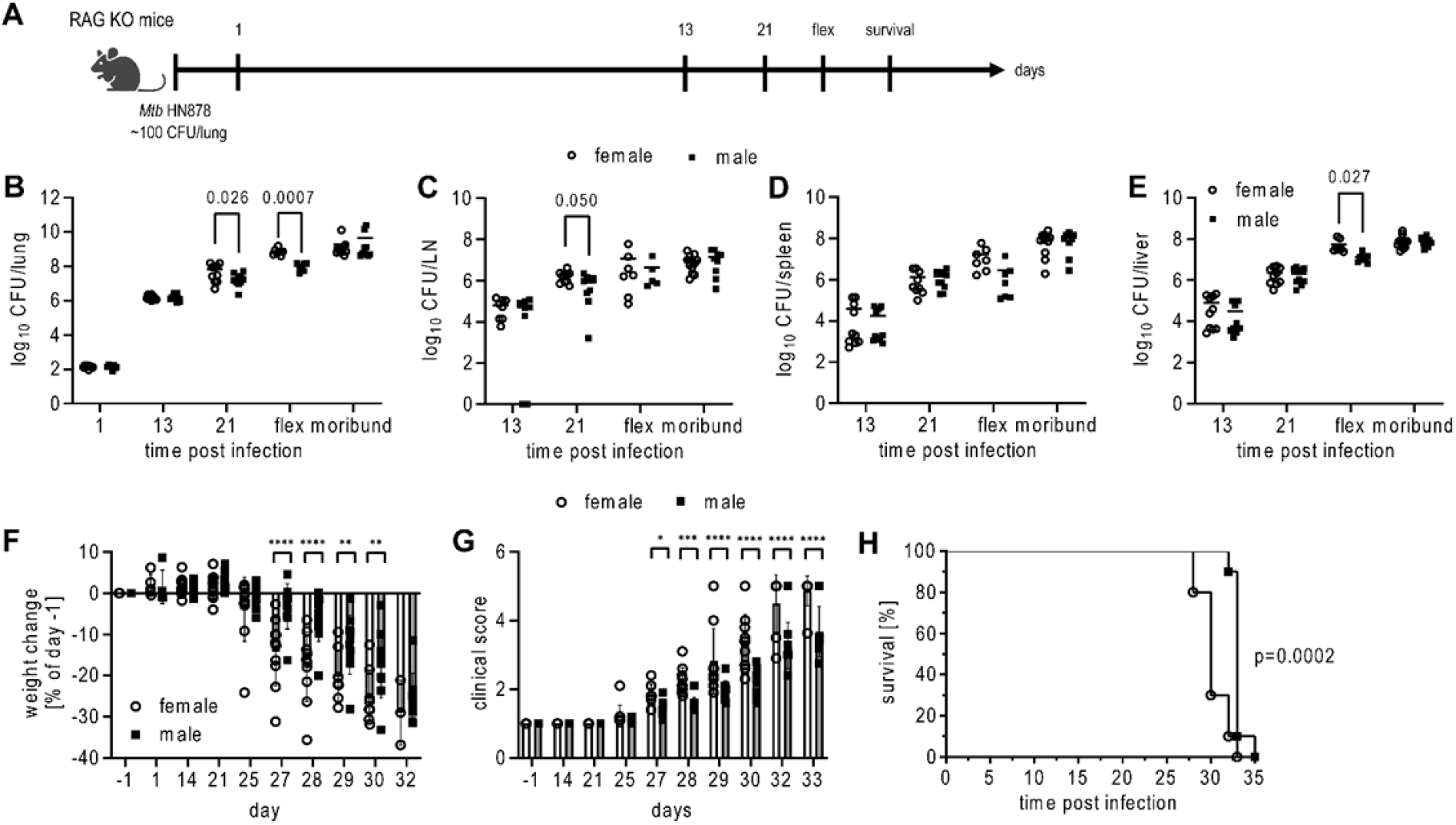
Increased susceptibility of female RAG KO mice to Mtb infection. A) Experimental setup. Female and male RAG KO mice were aerosol infected with *Mtb* HN878 and organs were collected at the indicated time points. Flex time point is defined as the time point where a mouse reached a score of 3. At the same time, a randomly selected mouse of the opposite sex, which had been assigned before the experiment, was also taken, regardless of its score. CFU of the lung (B), mediastinal LN (C), spleen (D) and liver (E) of female and male RAG KO mice at indicated time points. Body weight change (F), clinical score (G) and survival (H) of female and male RAG KO mice. B-E) Each data point represents one mouse from two experiments. Statistical analysis was performed by Student’s t-test (B-E), 2way ANOVA followed by Tukey’s multiple comparisons test (F+G) or log rank test (H).

While there were no differences in bacterial burden between the sexes on day 13, females exhibited significantly higher bacterial loads in the lungs and LN 21 days post-infection (Fig. 1B - 1E). As the disease progressed, evidenced by increased weight loss and clinical scores (Fig. 1F and 1G), bacterial loads rose further, especially in the lungs and livers of females, and were significantly higher than in males (Fig. 1B – 1E, flex). This aligns with the more pronounced weight loss and higher clinical scores observed in females compared to males (Fig. 1F and 1G). In moribund animals that had reached the endpoint according to our clinical scoring, the CFU did not differ between the sexes, likely reflecting comparable disease states. Overall, diminished bacterial control and accelerated disease progression culminated in earlier mortality in females (Fig. 1H). These findings contrast sharply with *Mtb* infection in immunocompetent C57BL/6 mice, where males are more susceptible than females ^5,6^.

### Sex differences in cytokine and chemokine production in response to Mtb in lungs of RAG KO mice

To examine the immunological environment in the lungs that was associated with differences in *Mtb* control in male and female RAG KO mice, we quantified the levels of inflammatory cytokines and chemokines. Fig. 2 illustrates sex-based differences in immune mediator profiles at day 21 post-infection and the later “flex” phase (no significant differences between sexes were observed on day 13 post *Mtb* infection; data not shown). Panels A and B present radar plots of cytokine and chemokine levels, comparing male and female RAG KO mice. Females exhibited elevated levels of several key inflammatory markers, including TNF, IL-1α, IL-1β, IL-6, IL-17A, CCL2, CCL4, CCL20, and CXCL1, as shown in Fig. 2C – 2K. These results suggest that female RAG KO mice have a generally heightened inflammatory response at both day 21 and during the flex phase, with statistically significant differences observed between sexes. However, CCL5 and IFN-γ stand out as notable exceptions to this trend. In contrast to the other immune mediators, both CCL5 and IFN-γ were significantly higher in males compared to females, particularly during the flex phase (Fig. 2L and 2M).

**Figure 2.**
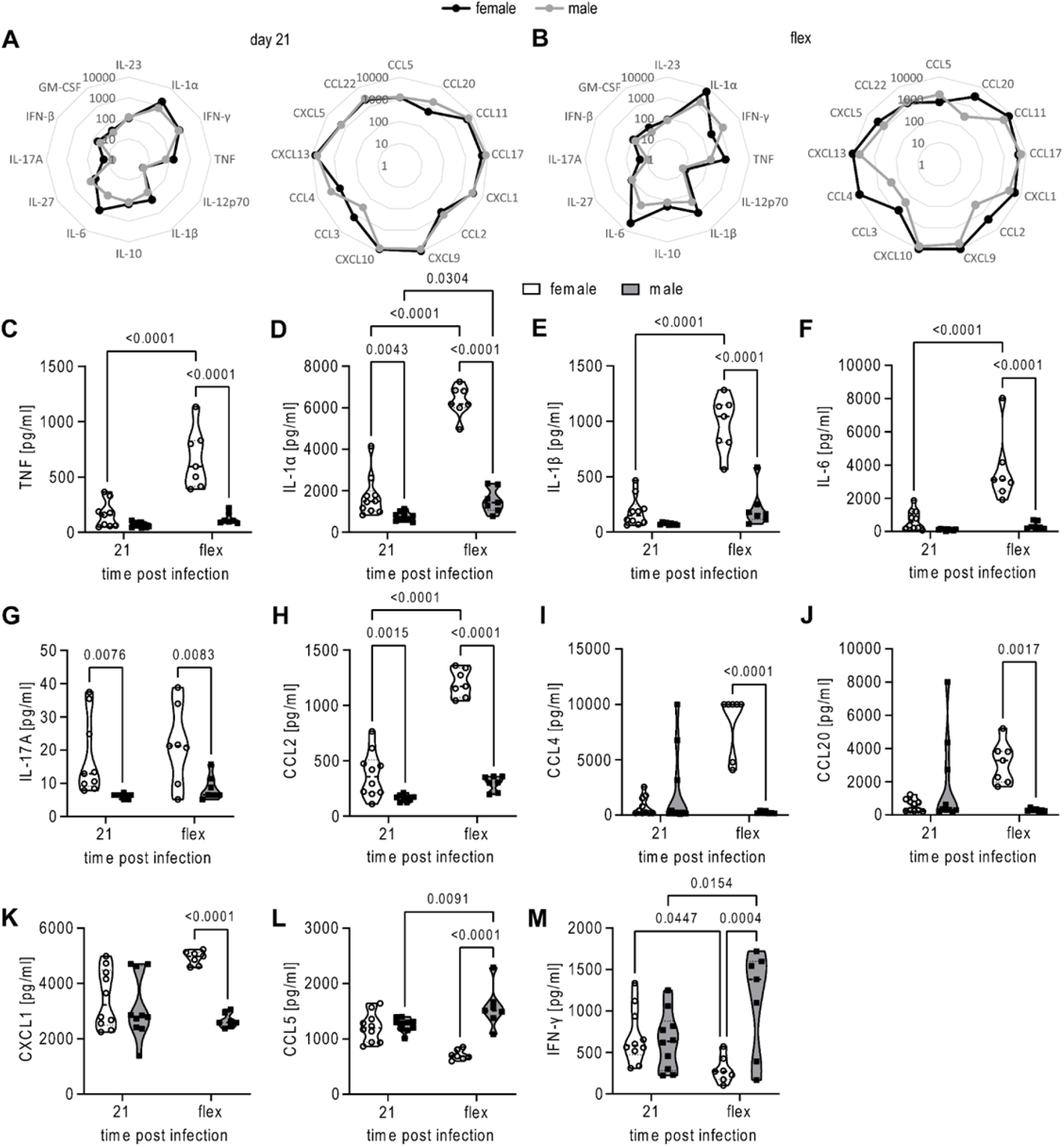
Inflammatory responses in the lungs of Mtb infected RAG KO mice. Female and male RAG KO mice were aerosol infected with Mtb HN878 and lungs were collected at the indicated time points. Radar chart of cytokines and chemokines measured in lung homogenates of female and male RAG KO mice at day 21 p.i. (A) and flexible time point p.i. (B). Concentrations of cytokines/chemokines are shown as pg/ml. C-M) Selected cytokines and chemokines measured in lung homogenates of female and male RAG KO mice at indicated time points. A + B) Data is represented as mean of cytokine and chemokine concentration from 2 experiments (A; n=10/group) or 1 experiment (B; n=7/group). C-M) Each data point represents one mouse from two experiments (d21; n=10/group) or one experiment (flex; n=7/group). Statistical analysis was performed by 2way ANOVA followed by Tukey’s multiple comparisons test.

These findings indicate marked differences between the sexes in the inflammatory response, driven by innate immunity. Since these cytokines and chemokines play a critical role in orchestrating immune cell recruitment, the data suggest that male and female RAG KO mice may exhibit distinct patterns of immune cell infiltration in the lung in response to *Mtb* infection.

### Females exhibit neutrophil-dominated inflammation, while males have more NK cells in the lungs

To support the cytokine and chemokine profiles, HE staining and immunohistochemistry (IHC) were conducted to evaluate lung pathology and cellular infiltration in *Mtb*-infected male and female RAG KO mice. The analysis revealed significantly larger areas of lung inflammation in females compared to males, particularly during the flex phase (Fig. 3A and 3B). This increased inflammation was associated with a higher abundance of CD68^+^ cells, which is a marker of macrophages and monocytes, in females than in males (Fig. 3C and 3D). Additionally, female lesions exhibited a significant influx of neutrophils, as evidenced by staining for 7/4^+^ cells (Fig. 3E and 3F). The elevated presence of neutrophils in female lungs corresponds with the increased levels of CCL20 and CXCL1 observed in females (Fig. 2J and 2K), both of which are known to play roles in neutrophil recruitment and activation. In contrast, NKR-P1C^+^ NK cells were more abundant in males than females (Fig. 3G and 3H). NK cells are a crucial early source of IFN-γ in the lungs during *Mtb* infection ^10^. Moreover, NK cell-derived IFN-γ was shown to regulate the pulmonary inflammatory response to *Mtb* in the absence of T lymphocytes by down-modulating neutrophil infiltration ^11^. Consistent with this, NKR-P1C^+^ cells were positively correlated with IFN-γ levels and negatively with 7/4^+^ cells and lung inflammation in our study (Fig. 3I – 3K). Conversely, the abundance of 7/4^+^ cells was positively correlated with the area of lung inflammation and clinical score (Fig. 3L and 3M).

**Figure 3.**
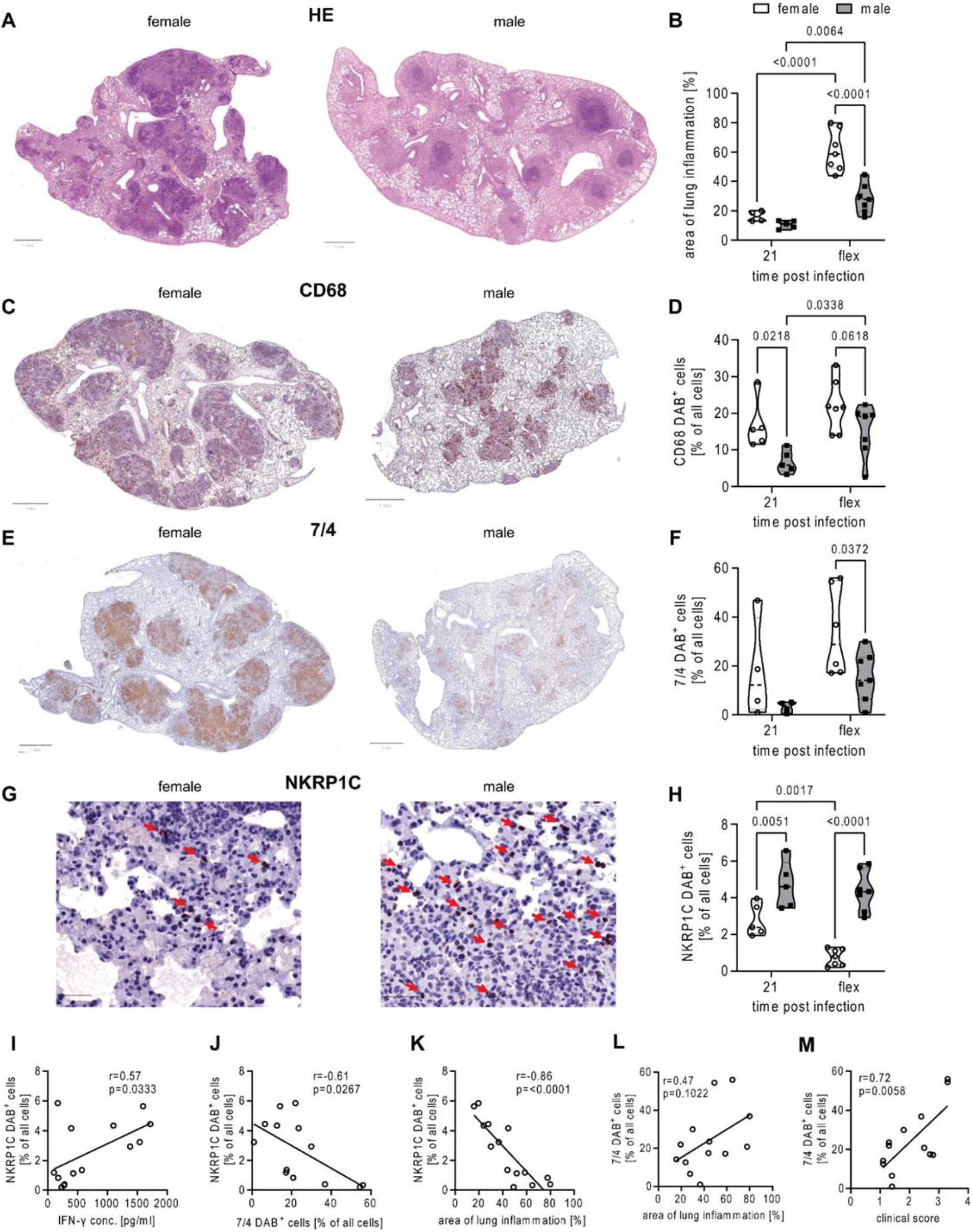
Increased NK cell numbers in male RAG KO mice associated with higher IFN-y responses and reduced neutrophil influx. Female and male RAG KO mice were aerosol infected with *Mtb* HN878. Lungs were collected at indicated time points and PFA-fixed, paraffin embedded tissue sections were stained with HE (A) or antibodies to detect (C) macrophages (CD68^+^), (E) neutrophils (7/4^+^), or (G) NK cells (NKR-P1C^+^). A, C, E, G) Representative micrographs from one female and male RAG KO mouse from the flexible time point p.i. are shown. Bar = 1 mm (A, C, E) or 50 µm (G). B, D, F, H) Quantitative analysis of the area of lung inflammation and respective immune cells as shown in (A, C, E and G). Correlation of NKR-P1C DAB^+^ cells with IFN-γ level (I), 7/4 DAB^+^ cells (J) and area of lung inflammation (K) as well as correlation of 7/4 DAB^+^ cells with area of lung inflammation (L) and clinical score (M) at the flexible time point p.i. B. D. F, H) Each data point represents one mouse from one representative experiment out of two (d21; n=5/group) or one experiment (flex; n=6-7/group). Statistical analysis was performed by 2way ANOVA followed by Tukey’s multiple comparisons test. I-M) Each data point represents one mouse from one experiment (flex; n=13-14). Correlation was calculated using Pearson correlation.

In conclusion, the data suggest that in the absence of adaptive immunity, female mice exhibit a neutrophil-dominated lung pathology, whereas males show a higher abundance of NK cells that may mitigate inflammation through IFN-γ production.

## Discussion

The initial immune response to *Mtb*, including the activation of innate immune cells and the subsequent inflammatory response, plays a crucial role in shaping the overall disease trajectory ^7,12^. In this study, we aimed to investigate whether early events in *Mtb* infection might already indicate the higher susceptibility and impaired long-term control observed in males ^5,6^. Our study reveals striking sex-specific differences in the innate immune response to *Mtb* infection in RAG KO mice, highlighting how the absence of adaptive immunity can lead to unexpected variations in disease susceptibility and progression. Contrary to what is observed in immunocompetent C57BL/6 mice, where males generally exhibit greater susceptibility to TB ^5,6^, female RAG KO mice demonstrated significantly poorer control over HN878 infection compared to males. This was evidenced by higher bacterial loads in the lungs, increased weight loss, and earlier mortality among females. The heightened susceptibility of female RAG KO mice was paralleled by an exaggerated inflammatory response, characterized by elevated levels of inflammatory cytokines and chemokines such as TNF, IL-1α, IL-1β, IL-6, IL-17A, CCL2, CCL20, and CXCL1. These factors are known to orchestrate immune cell recruitment and activation, and as a result, we observed a neutrophil-dominated lung pathology in females. This is in sharp contrast to males, who exhibited higher levels of NK cells and IFN-γ. In addition to its role in activating macrophages, IFN-γ has a regulatory role protecting from immunopathology by suppressing neutrophil recruitment ^13,14^. NK cells are an important early source of IFN-γ in the *Mtb* infected lung ^15^. Studies in immunocompetent C57BL/6 mice suggest that NK cells do not play a direct role in regulating TB immunity since depleting NK cells did not affect bacterial burden ^10^. However, NK cell-derived IFN-γ was indispensable for the survival of RAG deficient mice after *Mtb* infection where it limited neutrophil-driven immunopathology ^11^. Consistent with this, the increased abundance of NKR-P1C^+^ NK cells in males in our study correlated with reduced neutrophil infiltration and less severe lung inflammation, suggesting a protective role of NK cells in modulating inflammation in a sex-specific manner.

Sex differences in NK cells have been observed previously ^16-19^. All studies consistently showed that the number of NK cells was higher in males than in females, but the data on their activation status were inconsistent. Patin and colleagues found a larger number of activated NK cells in men than in women ^17^. Likewise, Huang and colleagues showed several pathways, such as the IFN-γ-mediated signaling pathway, and the positive regulation of immune effector processes to be up-regulated in male compared with female NK cells, which increased during aging ^16^. In contrast, the study from Cheng found enhanced effector function in female compared to male NK cells ^19^. These differences were primarily attributed to the X-linked epigenetic regulator UTX, encoded by the gene *Kdm6a*, which escapes X-inactivation ^20^. UTX influenced the expression of gene loci crucial for NK cell fitness and effector functions and was expressed at significantly lower levels in both male mouse and human NK cells ^19^. While these studies were conducted on immunocompetent mice, potential sex differences in the NK cell compartment of RAG KO mice remain unexplored. However, recent research has demonstrated that NK cells in RAG2-deficient mice are hyperactive ^21,22^. Both the number and frequency of NK cells were elevated in RAG2 KO mice compared to WT mice in peripheral blood, spleen, and bone marrow, with these cells also expressing higher levels of activation markers and IFN-γ. These hyperactive NK cells in RAG2-deficient mice showed enhanced antileukemia effect in several models of acute myeloid leukemia ^21^. Our study suggests that the potent NK cell response prevents neutrophil driven immunopathology in males but not in females. Currently the reasons for these sex differences remain unclear and deserve further evaluation.

Our findings reveal significant differences in disease outcomes following *Mtb* infection based on the presence or absence of adaptive immunity. In immunocompetent mice, males show greater susceptibility to *Mtb* infection ^4-6^, whereas in the absence of B and T cells, females exhibit heightened susceptibility. This shift underscores how adaptive immunity influences innate immunity and disease progression differently between sexes. Moreover, the pronounced neutrophil response in females and the potential regulatory role of NK cells in males underscore distinct innate mechanisms by which each sex responds to *Mtb* infection. Understanding these immune differences, and how innate and adaptive immunity interact in males and females, is crucial for designing interventions that address the unique immune profiles of each sex and enhance overall disease management.

## Acknowledgements

We would like to thank Silvia Maass for culturing *Mtb*; the staff of the animal facility at the Research Center Borstel for animal care; Christian Rosero and Jasmin Tiebach for their help with the immunohistochemistry. This work was supported by in-house funding from the Research Center Borstel.

## Author contributions

B.E.S. and D.H. conceived and designed the experiments; D.H., L.E. and L.v.B. performed the experimental work; D.H. analyzed the data; T.G. and H.L. provided advice, reagents and mice; B.E.S. and D.H. wrote the paper. All authors revised the manuscript.

